# Focal persistence and phylodynamics of Heartland virus in Georgia

**DOI:** 10.1101/2024.10.07.616953

**Authors:** Stephanie Bellman, Nima Shariatzadeh, Tyshawn Ferrell, Audrey Long, Leah Aeschleman, Ellie Fausett, Tim Walsh, Seana Cleary, Isabella Roeske, Erick Ojeda, Madison Schwab, Hannah Dakanay, Sam R Telford, Heidi K Goethert, Gonzalo Vazquez-Prokopec, Anne Piantadosi

**Affiliations:** Gangarosa Department of Environmental Health, Rollins School of Public Health, Emory University, Atlanta, GA, US; Department of Environmental Sciences, Emory University, Atlanta, GA, US; Department of Pathology and Laboratory Medicine, Emory University, Atlanta, GA, US; Department of Microbiology and Immunology, Emory University, Atlanta, GA, US; Graduate Program in Biochemistry, Cell, and Developmental Biology, Emory University, Atlanta, GA, US; Graduate Program in Population Biology, Ecology, and Evolution, Emory University, Atlanta, GA, US; Department of Infectious Disease and Global Health, Cumming School of Veterinary Medicine, Tufts University, Grafton, MA, US; Department of Medicine, Division of Infectious Diseases, Emory University, Atlanta, GA, US

## Abstract

Heartland virus (HRTV) is an emerging tick-bone virus associated with severe illness in the U.S. There are large gaps in knowledge of HRTV diversity, evolution, and transmission due to a paucity of HRTV-positive samples and genome sequences. We identified a focal site of HRTV- positive *Amblyomma americanum* ticks in central Georgia and developed a novel multiplex- amplicon sequencing assay to generate full HRTV genome sequences. By screening over 21,000 field-collected ticks from 2021-2023, we identified six positive pools. Five were collected from the site in central Georgia where our group first detected HRTV-positive ticks in 2019, and one from a site in western Georgia approximately 175 km away. The HRTV genome sequences from Georgia were highly related, even across this distance and over five years. Reference HRTV genome sequences from across the U.S. were also geographically clustered. Time-scaled phylogenetic analysis suggested recent spread of HRTV in the U.S., with all available sequences sharing a common ancestor within the last 300 years, and sequences from Georgia sharing a common ancestor within the last 40 years. Our observed spatial clustering of HRTV and the high degree of genetic conservation in our persistent focus suggest the importance of small spatial dynamics in HRTV transmission ecology.

**Author Summary:** Heartland virus (HRTV) was first discovered in humans in 2009 and has since caused over 60 cases of severe and fatal disease in the United States. HRTV is transmitted by the lone star tick, *Amblyomma americanum*, across the Southeast, East coast, and Midwest. Little information is known about how this virus circulates and changes across time and space due to a lack of genetic data. Here, we created a new procedure to generate more genetic sequence data for HRTV and collected over 21,000 ticks to screen for HRTV across three years in Georgia. We generated 6 new HRTV sequences and compared them to existing sequences from our group in Georgia, and across the country, finding evidence of regional clustering of HRTV and highly related HRTV across time in Georgia. Our analyses additionally found that this virus was likely introduced to the U.S. in the last 300 years. Our study provides new context and information in understanding the landscape and transmission of HRTV in the U.S.

## Introduction

In 2009, two farmers in northwestern Missouri presented with symptoms similar to ehrlichiosis and were found to have a novel pathogen, Heartland virus (HRTV).(1) Originally categorized as a *Phlebovirus*, HRTV was recategorized to the genus *Bandavirus* in the family *Phenuiviridae,* order *Bunyavirales* in 2020.(2) HRTV is vector-borne and transmitted by the lone star tick (*Amblyomma americanum*), whose range spans the majority of the eastern half of the United States.(3, 4) To date, HRTV has been associated with over sixty cases of human disease in fourteen U.S. states and has an observed case fatality rate of 5-10%.(5) HRTV, like other bunyaviruses, has a segmented negative-sense single-stranded RNA genome with three segments.(1) The L segment encodes the RNA-dependent RNA polymerase (RdRp) and is 6.4 kb long. The M segment encodes two viral glycoproteins (Gn, Gc) and is 3.4 kb long. Lastly, the S segment employs an ambisense strategy to encode the nucleoprotein (Np) and nonstructural protein (NSs) in 1.7 kb. HRTV is most closely related to Dabie bandavirus (DBV) (previously known as Severe Fever and Thrombocytopenia Syndrome virus, SFTSV) which circulates in China, Japan, South Korea, Vietnam and Taiwan.(6) While different genotypes of DBV have been documented across Asia(7) and have been associated with different pathogenicity in humans,(8) very little is known about the genotypic and phenotypic diversity of HRTV. This knowledge gap stems from limited surveillance, low HRTV prevalence in ticks, (9–11) and few detected human cases.(12) HRTV has been identified in *A. americanum* populations in eight states across the U.S., spanning the Northeast, Southeast, and Midwest.(13) Prior surveillance studies for HRTV in ticks have been episodic, typically responding to newly discovered human cases, and continuing for one to two years.(9–11, 14) Few studies have continued surveillance beyond two years, and there is very little HRTV genome sequence data available, creating a gap in understanding the enzootic transmission and evolutionary dynamics of HRTV.

The objective of this study was to begin to fill this knowledge gap through long term surveillance at established HRTV-positive tick sites in central Georgia (GA), where our group initially detected HRTV in 2019.(15) In our prior study, we found a minimum infection rate of 0.46/1000 HRTV-positive *A. americanum* ticks at two locations where HRTV seropositivity was previously reported in deer populations,(16) and adjacent to the county where a human case was reported.(12) In the present study, we build upon this by performing extended HRTV surveillance in GA across multiple years. We use a novel multiplex amplicon sequencing approach to generate full HRTV genome sequences and analyze phylogenetic and phylogeographic relationships of HRTV in GA and across the U.S., gaining insight into potential mechanisms of virus transmission and evolution.

## Methods

### Field collections

Ticks were collected from March-August 2021-2023, corresponding to the seasonality of *Amblyomma americanum* in Georgia.(17) Collection locations varied by year: in 2021, repeated sampling was conducted at the two sites where our group detected Heartland virus in 2019(15); in 2022, sampling was performed in forty-three locations across the state(18); and in 2023, five locations with the highest density of *A. americanum* from the previous years were repeatedly sampled to maximize potential HRTV detection. Free flagging was conducted at each site according to previously described protocols,(19) with the addition of transect sampling during 2022.(18) Ticks were transported live in plastic vials to the laboratory and microscopically identified with taxonomic keys.(20, 21)

### Tick processing, RNA extraction, and qRT-PCR

Surface disinfection was conducted on ticks in field vials using a wash of 10% bleach for five minutes, 3% hydrogen peroxide for five minutes, and three rinses of distilled water for five minutes each. The ticks were pooled in groups of approximately five adults or twenty-five nymphs per species per site. Differing amounts of BA-1 diluent (1× medium 199 with Hanks balanced salt solution, 0.05 mole/L Tris buffer [pH 7.6], 1% bovine serum albumin, 0.35 g sodium bicarbonate/L, 100 μg/L streptomycin, 1 μg/mL amphotericin B) were added to each pool based on the number of ticks in the pool (1 ml for >1 adult or >10 nymphs, 0.5 mL for 1 adult or <10 nymphs) to preserve concentrations of the ground ticks in media. Pools were ground thoroughly using a 7-mL glass TenBroeck grinder (Fisher Scientific) with alundum bedding material (Fisher Scientific) as an abrasive.

Tick homogenates underwent RNA extraction (QIAmp TNA Extraction Kit) following the manufacturer’s protocol (Supplemental Table S1). Extracted RNA was tested using quantitative reverse-transcriptase PCR (qRT-PCR) with primer-probe set 1 for the S segment of HRTV described by Savage et al.(10) The AgPath-ID One-Step RT-PCR kit (Thermo Fisher Scientific) was used as per the manufacturer’s instructions, with cycling conditions of 45°C for 10 minutes, 95°C for 10 minutes, 45 cycles of 95°C for 15 seconds and 60°C for 45 seconds. The reaction mix and primer and probe sequences are available in Supplemental Table S2. Samples were tested in duplicate with positive controls (extracted HRTV virus stock from Romer et al.(15)) and negative controls (water) on each plate. As a quality control check for RNA extraction, tick actin RT-qPCR was run in parallel using the Applied Biosystems Power SYBR Green RNA-to-Ct 1-step PCR Kit (Fisher Scientific) with primers, probes, and reaction conditions listed in Supplemental Table S3.

### Infection rate estimation

The minimum infection rate (MIR) of HRTV was estimated by site and year by dividing the number of positive pools at a location by the number of *A. americanum* of the same life stage collected at the site and year and multiplying by 1,000.

### HRTV multiplex amplicon sequencing primer design

Multiplex amplicon sequencing primers for HRTV were designed with primalscheme (v1.4.1)(22) using an alignment of 4 (L) or 5 (M and S) complete reference HRTV sequences.(23) Primers that had high annealing temperatures (65°C or greater) or had SNPs expected to affect primer binding, as determined by comparison to additional partial genome sequences, were redesigned in Geneious Prime (v2022.2). A set of 36 L, 18 M, and 10 S primer pairs were created to cover the entire genome, with each primer pair amplifying an approximately 250bp region (Supplemental Table S4). The HRTV-specific primer sequences were concatenated with a universal Nextera adapter sequence to allow Nextera XT indexing and amplification.(24, 25) Primers were synthesized by IDT, pooled, and tested using a previously-sequenced HRTV-positive sample.(15) Primer concentrations were optimized to ensure approximately equal coverage across the entire genome.

### HRTV full genome multiplex amplicon sequencing

RNA samples that tested positive for HRTV via RT-qPCR underwent heat-labile dsDNase treatment (ArticZymes), single-stranded cDNA synthesis (Fisher/Invitrogen) and multiplex PCR using Q5 Hot Start High-Fidelity Polymerase (New England Biolabs) using the custom-designed primers. The resulting amplicons were indexed using Nextera XT (Illumina) and quantified using the KAPA Universal Complete Kit (Roche). Libraries were pooled to equimolar concentration and sequenced with 300bp paired-end reads on an Illumina MiSeq. Reads were filtered for quality, and adaptor sequences were removed, before undergoing reference-based assembly using ViralRecon v. 2.6.0 with reference sequences L: NC_024495, M: NC_024494, and S: NC_024496. A more complete protocol for this sequencing method can be found in Supplemental Text S1.

### Phylogenetic analysis

Twelve reference sequences of Heartland virus L, M, and S segments present in GenBank as of March 1^st^, 2024 were downloaded for analysis. Selection criteria included: related to a field collected case (tick or human), no identical isolates (if an isolate was sequenced more than once, the earliest was selected), and >95% genome coverage. Newly-generated sequences and reference sequences were aligned by segment in Geneious Prime (v2022.2.2) using MAFFT (v7.450) with default parameters.(26, 27) Maximum-likelihood phylogenetic trees were generated for each segment using PhyML 3.0,(28, 29) with best fit substitution models: GTR+G for the L segment, HKY85+G for the M segment, and K80+I for the S segment. Trees were subsequently visualized and annotated using iTOL(30). Reassortment was assessed through two methods 1) by visually comparing segment relationships across different clades within the trees and 2) by testing aligned concatenated coding regions with the recombination detection program, RDP4.(31)

### Bayesian time-scaled phylogenetic analysis

To construct time-scaled phylogenies, the temporal signal of each segment’s coding region was evaluated using TempEST(32), and one outlier was removed using a threshold of 0.03 with unbalanced residuals on the residual histogram distribution (reference sequence OK480062.1, Supplemental Figure S1). There was evidence for moderate temporal signal that varied by segment, with root-to-tip correlation coefficients of 0.59 for the L segment, 0.55 for the M segment, and 0.33 for the S segment. Time-scaled phylogenies were therefore constructed only for the L and M segments, using BEAST 2.7.6 with a chain length of 600 million.(33) We used a GTR+gamma nucleotide substitution model, a constant coalescent population, and a strict molecular clock with a prior of 2E-4,(34–36) with a lognormal distribution. The log files were examined with Tracer v1.7.2 and confirmed to have ESS values >200 for all parameters.(37) Maximum clade credibility trees were generated with 10% burn-in using TreeAnnotator v2.7.6.(33) More complex models, including a relaxed molecular clock and skyline models of population growth, did not converge, likely as a result of the limited HRTV data available.

### Geographic clustering analysis

The Mantel test(38) was used to assess the relationship between geographic distance and genetic divergence. Because exact coordinates were not known for the reference sequences, the location was estimated using the finest resolution information present. For samples with only a state recorded for the source location in GenBank, the center point of the most central county in the state was assigned, using USGS(39) and Latitude.to (v1.64-im). For samples with a county and a state recorded for the source location in GenBank, the center of the county listed for that sequence was assigned. For sequences associated with publications, the smallest resolution estimate was made based on recorded location information.(9, 11, 40) Lastly, for the two initial Missouri human case sequences, Andrews county and Nodaway county centers were used based on Savage et al.(40) The Mantel test was run using the vegan package (v2.5-7) in R (v4.0.4) using 1000 permutations.

### Host bloodmeal analysis

Nucleic acid extracted from HRTV-positive *A. americanum* pools and matched HRTV-negative *A. americanum* pools – containing a similar number of ticks, and the same life stage, collection site, and collection date – were tested by multiplex qPCR targeting retrotransposons of potential hosts, according to established protocols.(41, 42) Species included in the assay were: mice, voles, rabbits, birds, shrews, squirrels/chipmunks, deer, skunks and opossums.

## Results

### Six HRTV positive tick pools were detected across central and western Georgia, U.S., from 2021 to 2023

In total, 21,386 ticks were collected across the three year period: 7,687 in 2021, 3,444 in 2022, and 10,255 in 2023 (Table 1). Most (20,959, 98%) were *A. americanum* adults or nymphs. Six pools of *A. americanum* ticks tested positive for Heartland virus: 1 pool of 19 nymphs from 2021, 2 pools of 27 nymphs and 28 nymphs from 2022, and 3 pools of 25 nymphs, 5 adult females, and 25 nymphs in 2023 (Table 2). Five of the six HRTV-positive pools were collected at ST road site (ST) in central GA, and the sixth was collected at Chattahoochee Bend State Park (CB) in 2023 (Figure 1). The minimum infection rates (MIRs) calculated for each site and year were 0.33 per 1000 nymphs at ST in 2021, 1.97 per 1000 nymphs at ST in 2022, 0.59 per 1000 nymphs at ST in 2023, 2.56 per 1000 adults at ST in 2023 and 0.46 per 1000 nymphs at CB in 2023.

**Figure 1.**
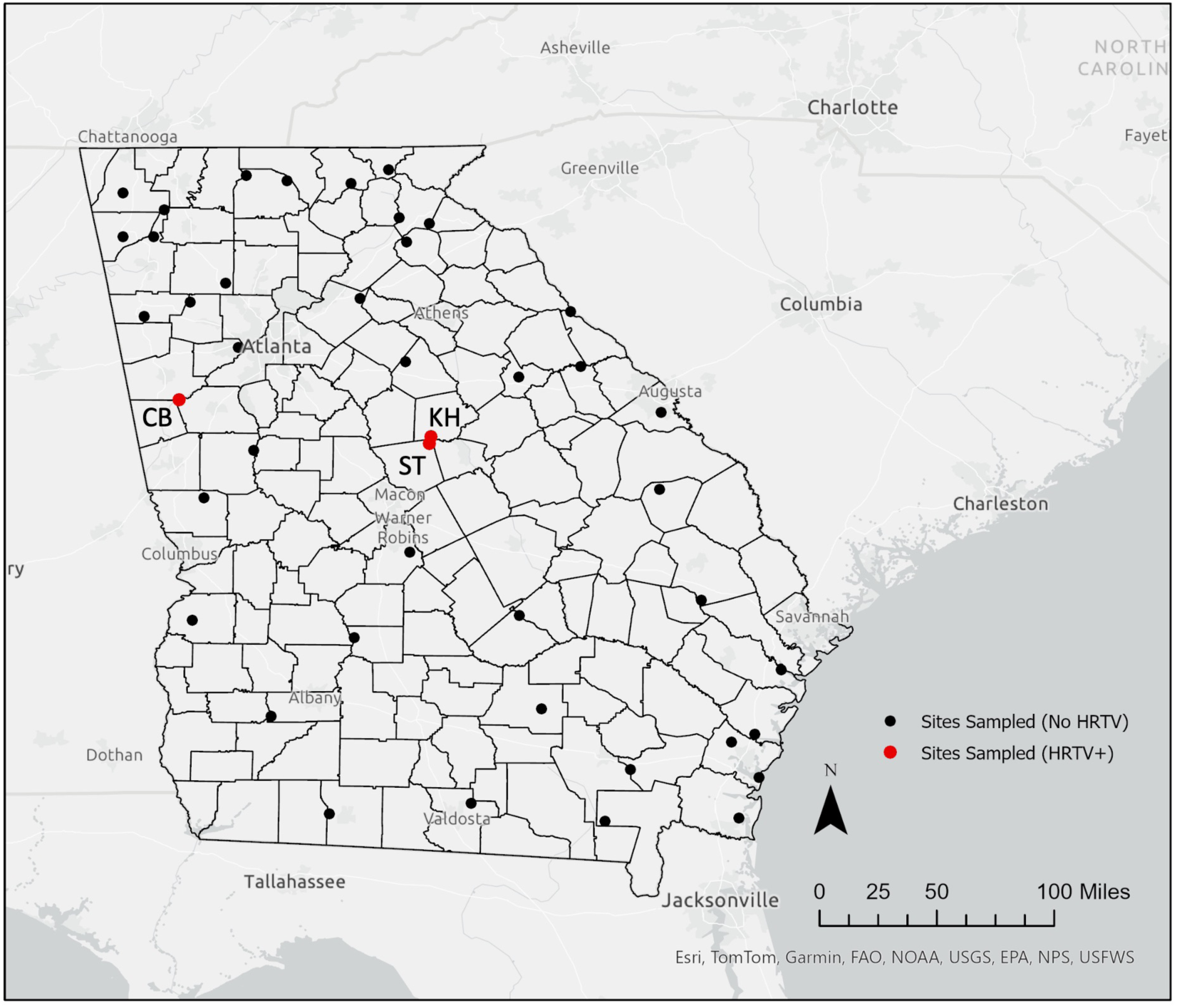
**Map of collection sites visited from 2021-2023**. State and county boundaries for Georgia are highlighted in black. Black dots represent locations sampled where HRTV positive pools of ticks were not found. Red dots represent locations where HRTV positive pools were found with each location labeled with its two letter site code (KH = KH road site, ST = ST road site, CB = Chattahoochee Bend State Park). Though no positive pools of HRTV were found from 2021- 2023, KH is still highlighted due to presence of positive ticks from 2019 detected by our group as originally documented in Romer et al(15) and described in Table 2. Map created in ArcGIS Pro v3.1.0.

**Table 1.**
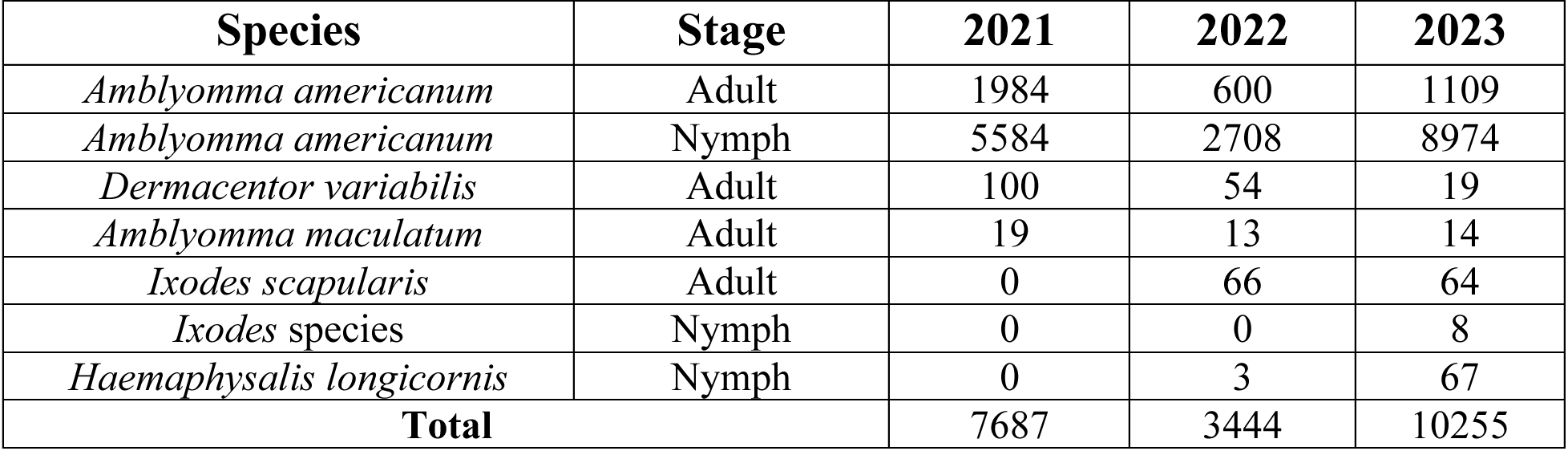
Species and life stage of ticks collected during field sampling from 2021-2023.

**Table 2.** Heartland positive pool characteristics.

| Year | Lab Code | Location | Date Collected | Number of Ticks | Life Stage |
| --- | --- | --- | --- | --- | --- |
| 2019* | 23 | KH | 4/28/19 | 5 | Adult (Female) |
| 2019* | 26 | KH | 4/28/19 | 25 | Nymph |
| 2019* | 504 | ST | 6/14/19 | 5 | Adult (Male) |
| 2021 | 404 | ST | 6/15/21 | 19 | Nymph |
| 2022 | 100 | ST | 5/12/22 | 27 | Nymph |
| 2022 | 187 | ST | 6/10/22 | 28 | Nymph |
| 2023 | 56F | ST | 5/12/23 | 25 | Nymph |
| 2023 | 432R | ST | 6/16/23 | 5 | Adult (Female) |
| 2023 | 642T | CB | 7/19/23 | 25 | Nymph |

All pools consist of *Amblyomma americanum*. Location abbreviations: KH = KH road site, ST = ST road site, CB = Chattahoochee Bend State Park. \*2019 HRTV-positive pools were originally published in Romer et al.(15)

### HRTV sequences from GA are highly related across space and time

Full HRTV genome sequences were generated using our novel HRTV-specific multiplex amplicon protocol, with 97-100% coverage achieved across all three segments from the six positive tick pools. Including the three HRTV positive tick pools collected and sequenced by our group in 2019 from sites ST and KH,(15) the nine GA sequences were highly similar with 99.52- 99.99% nucleotide identity across the entire genome and similar identity relationships across the individual segments (Table 3). The largest genetic differences were seen between the sample collected at site CB in 2023 (Sample 642T) and other samples from GA, collected approximately 175 km away. Sample 642T differed from the other GA sequences by 44-51 single nucleotide polymorphisms (SNPs), while the sequences from sites KH and ST differed from one another by only 1-27 SNPs (Table 3). Few non-synonymous SNPs were found in sample 642T: 4 (of 24 total SNPs) in the L segment, 1 (of 13) in the M segment, and none in the S segment. Interestingly, while three of these non-synonymous SNPs were unique to sample 642T, two SNPs in the L segment (T168A and R183K) were shared with the other twelve reference sequences obtained from across the country.

**Table 3.** Pairwise genetic distances between HRTV sequences in Georgia.

| Location |  | Site KH |  | Site ST |  |  |  |  |  |
| --- | --- | --- | --- | --- | --- | --- | --- | --- | --- |
| Site KH |  | HRTV-23_T_GA_KH_<br>2019 | HRTV-26_T_GA_KH_<br>2019 | HRTV-100_T_GA_ST_<br>2022 | HRTV-404_T_GA_ST_<br>2021 | HRTV-187_T_GA_ST_<br>2022 | HRTV-504_T_GA_ST_<br>2019 | HRTV-056F_T_GA_ST_<br>2023 | HRTV-432R_T_GA_ST_<br>2023 |
|  | HRTV-26_T_GA_KH_<br>2019 | 3 (99.97%) |  |  |  |  |  |  |  |
| Site ST | HRTV-100_T_GA_ST_<br>2022 | 10 (99.91%) | 7 (99.93%) |  |  |  |  |  |  |
|  | HRTV-404_T_GA_ST_<br>2021 | 8 (99.92%) | 7 (99.93%) | 2 (99.98%) |  |  |  |  |  |
|  | HRTV-187_T_GA_ST_<br>2022 | 11 (99.9%) | 8 (99.92%) | 1 (99.99%) | 3 (99.97%) |  |  |  |  |
|  | HRTV-504_T_GA_ST_<br>2019 | 23 (99.78%) | 22 (99.79%) | 23 (99.78%) | 21 (99.8%) | 24 (99.77%) |  |  |  |
|  | HRTV-056F_T_GA_ST_<br>2023 | 11 (99.9%) | 10 (99.91%) | 7 (99.93%) | 7 (99.93%) | 8 (99.92%) | 26 (99.75%) |  |  |
|  | HRTV-432R_T_GA_ST_<br>2023 | 12 (99.89%) | 11 (99.9%) | 8 (99.92%) | 10 (99.91%) | 9 (99.91%) | 27 (99.74%) | 7 (99.93%) |  |
| Site CB | HRTV-642T_T_GA_CB_<br>2023 | 47 (99.56%) | 44 (99.58%) | 46 (99.56%) | 46 (99.56%) | 47 (99.56%) | 51 (99.52%) | 47 (99.56%) | 46 (99.56%) |

Number of SNPs and percent nucleotide identity is displayed across the concatenated genome of the three segments excluding gaps. Background color gradient represents nucleotide identity with darker colors representing higher nucleotide identity. Sites are delineated by bolded lines.

When comparing all twenty-one HRTV sequences from the U.S., the nucleotide identity ranged from 92.76-99.99% across the three segments, equating to between 1 and 782 SNP differences (Supplemental Table S5). Across the HRTV genome, the GA sequences shared 70 SNPs (35 in L, 20 in M, 12 in S), 6 of which (1 in L, 2 in M, 3 in S) were non-synonymous. While the GA sequences were all derived from tick pools, the twelve available reference sequences from GenBank were from both tick pools (5) and human infections (7). There were no SNPs common to HRTV sequences from only ticks or only humans.

### U.S. HRTV sequences demonstrate geographic clustering and potential reassortment

The 21 U.S. HRTV sequences generally clustered geographically on maximum-likelihood phylogenetic trees for each segment (Figure 2). Across all trees, the GA sequences clustered together and the sample collected at western site CB (HRTV 642T) was the most divergent. The two sequences from NY ticks similarly clustered together across all segments. Furthermore, samples from neighboring states generally clustered together (OK/KS/MO and TN/KY) in each segment, regardless of date of sample collection or host. The Mantel test also supported geographic clustering, with a correlation of 0.53 (p-value=0.001) between genetic distance and spatial distance.

**Figure 2:**
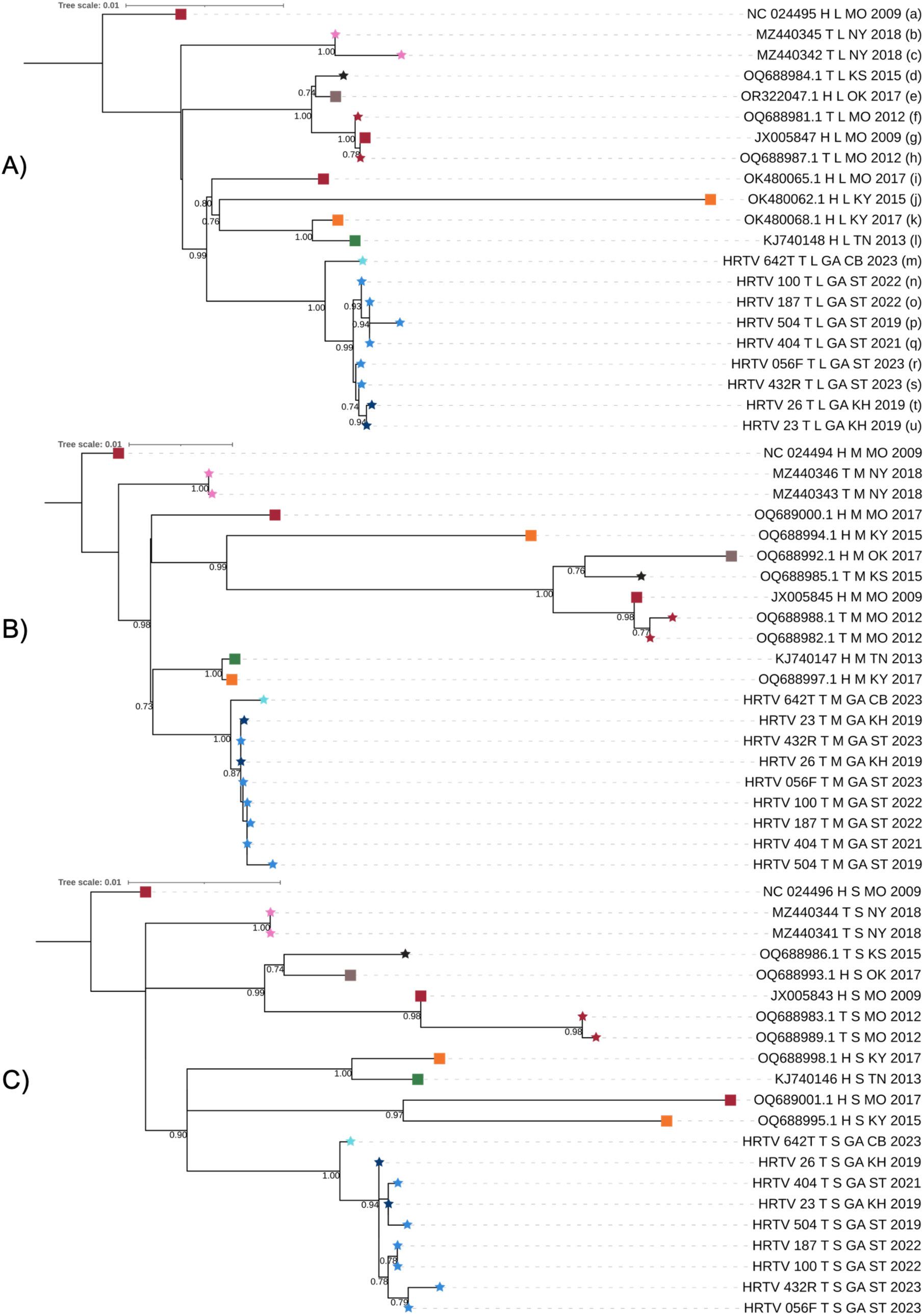
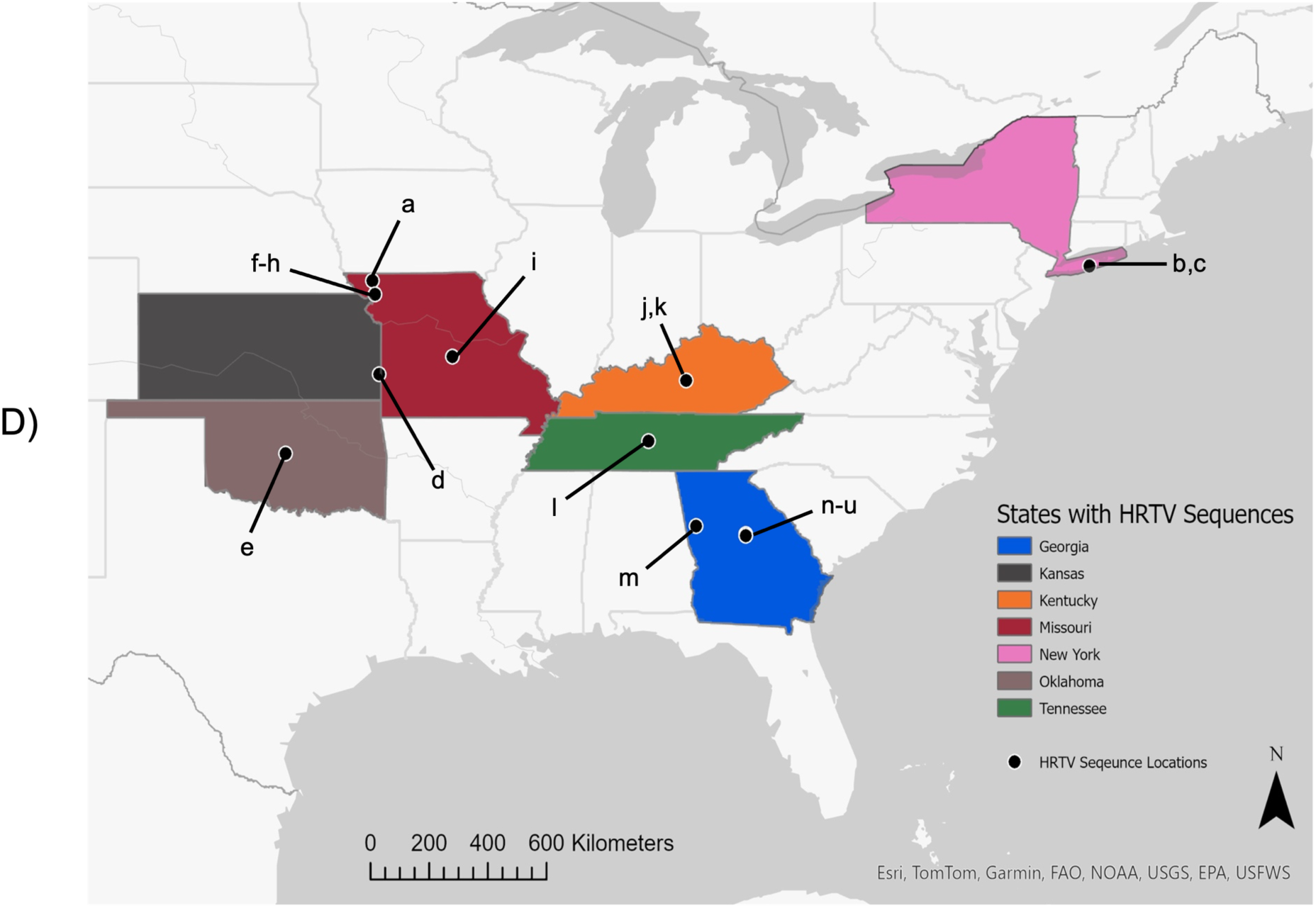
Phylogenetic and geographic relationships of HRTV sequences in the U.S. Segments were aligned separately, and maximum likelihood trees are displayed for the L segment (A), M segment (B), and S segment (C). Reference sequences from GenBank were named by the accession code followed by T (tick sample) or H (human sample), the segment, the state abbreviation for the collection location, and the year of collection. Sequences generated in this study were named by HRTV, sample number, T, GA, collection site, and year collected. All trees were rooted on the reference sequence from MO Patient 1. Ultrafast bootstrap values above 0.7 are displayed on the tree. Tips are labeled with a star for tick samples and square for human samples. The tip color indicates the location of collection. D) A map of the Eastern United States with states colored as in the phylogenetic trees (panels A-C). Locations for each sequence included are marked with a black dot. Dots are placed where either 1) sequences locations were documented in the literature(9, 11, 40) or 2) in the center of the state/county reported when finer resolution location data was unavailable. Dots are additionally marked with a-u and correspond to the sequences in the tree marked in panel A. Map created in ArcGIS Pro v3.1.0.

While the overall phylogenetic structure was similar across segments, we observed two examples of potential reassortment between segments. First, sequences OQ688994-5, collected from a human case in KY in 2015, clustered with the other KY and TN sequences in the L segment but with the MO, OK, and KS clade in the M segment, and with a single MO sequence in the S segment, all with moderate to high ultrafast bootstrap support (0.76, 0.99, 0.97 respectively). The second example of potential reassortment is from our sample 432R, collected at site ST in 2023. It clusters with the site KH sequences from 2019 (23 and 26) in the L segment, but with the other site ST sequences (187, 100, 056F) in the S segment, with moderate ultrafast bootstrap support (0.74, 0.79). These differences in tree topology may indicate reassortment, as has been reported for the related Dabie bandavirus. RDP4 was also used to assess for reassortment events between the HRTV segments and did not identify any events with high certainty.

### Contemporary HRTV strains in the U.S. emerged within the last 300 years

Time-scaled Bayesian phylogenetic analysis showed an evolutionary rate of 1.1 x10^-4^ substitutions/site/year (s/s/y) in the L segment and 1.8 x10^-4^ s/s/y in the M segment; this was not performed for the S segment given the lower temporal signal observed. Based on the L segment, the time to most recent common ancestor (tMRCA) for the GA clade was 23 years ago (95% HPD 12-42); the tMRCA for the GA clade and its nearest neighbor, the MO/KY/TN clade, was 85 years ago (95% HPD 46-153); and the overall root of the tree was 105 years ago (95% HPD 57-187) (Figure 3). Based on the M segment, the tMRCA for the GA clade was 16 years ago (95% HPD 8-29); the tMRCA for the GA clade and its nearest neighbor, the KY/TN clade, was 56 years ago (95% HPD 31-99); and the overall root of the tree was 150 years ago (95% HPD 78-286) (Figure 3).

**Figure 3:**
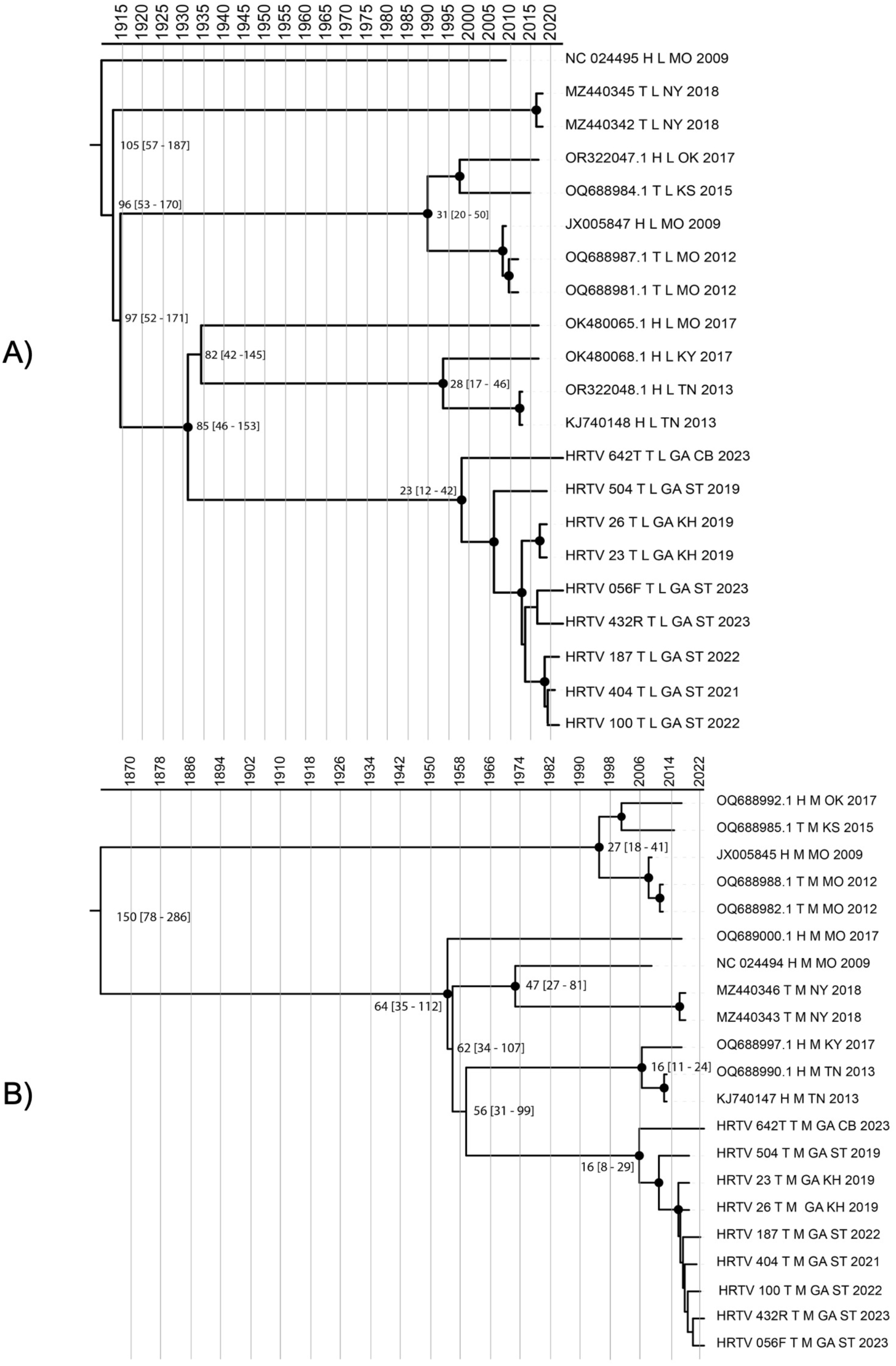
Time-scaled Bayesian phylogenetic tree of HRTV segments L (A) and M (B). Nodes are labeled with median time to recent common ancestor (tMRCA) with [95% HPD].

### *Amblyomma americanum* ticks feed on a variety of wildlife in Georgia

To begin to evaluate potential reservoir species for HRTV, a total of 12 *A. americanum* pools from GA underwent host bloodmeal analysis using retrotransposon qPCR. Each HRTV-positive pool was matched with an HRTV-negative pool with ticks of the same life stage, collection site, and collection date (Table 4) . The HRTV-positive pools of nymphs from ST (samples 404, 187, 100, and 56F) tested positive for birds, deer, rabbits, and squirrels/chipmunks. Their matched controls from HRTV-negative pools had similar results, with two exceptions: HRTV-negative pool 403 was positive for rabbit while its matched HRTV-positive pool 404 was not, and HRTV- positive pool 56F was positive for squirrel/chipmunk, while its matched HRTV-negative pool 55E was not. Both pools of adult ticks from site ST tested negative for most hosts, other than rabbit in the HRTV-negative pool. Finally, the nymph pools from CB both tested positive for deer, and the HRTV-negative pool additionally tested positive for squirrels/chipmunks.

**Table 4.**
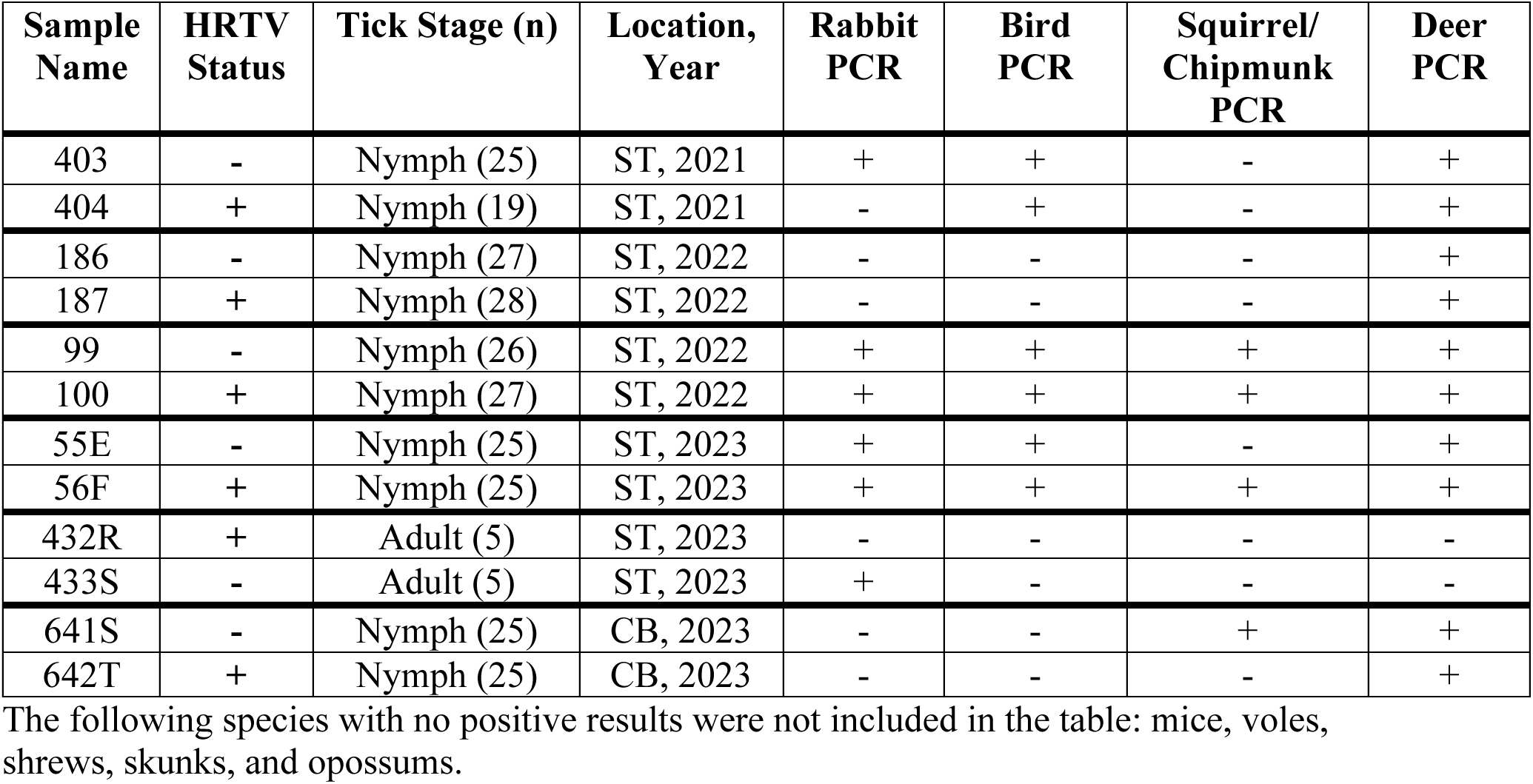
Host Bloodmeal PCR results.

## Discussion

We identified a persistent focus of genetically conserved HRTV in central Georgia. We collected over 21,000 ticks across three years and found minimum infection rates (MIRs) ranging from 0.33 per 1000 nymphs in 2021 to 2.56 per 1000 adults in 2023, compatible with our prior work(15) and other studies.(40, 43) Using a novel multiplex amplicon sequencing assay, we generated six new HRTV genome sequences and examined phylogenetic relationships across multiple years and locations.

One key finding was the very high genetic conservation of HRTV from ticks collected at a focal site in central GA across five years. Studies of Powassan virus, an unrelated tick-borne virus also endemic to the U.S., have similarly reported foci of genetically conserved viruses that were stable across multiple years, but have also reported geographic dispersal of closely-related POWV strains.(44–46) We did not observe geographic dispersal of closely-related HRTV strains, instead we observed phylogenetic clustering by location throughout the U.S., though it is notable that sequence data are limited, with only 12 reference sequences available from elsewhere in the U.S. and our 9 from Georgia.

A second key finding was that time-scaled phylogenetic analysis indicated recent spread of HRTV in the U.S., with all 21 sequences sharing a common ancestor within the last 300 years, and sequences from Georgia sharing a common ancestor within the last 40 years. This timeline is compatible with historical patterns of *A. americanum* prevalence.(47) In the 18^th^ century, dense forest supported high populations of deer and associated ticks, however deforestation and decimation of deer populations in the 19^th^ century led to large-scale *A. americanum* range retraction. The resulting fracturing of tick populations could have generated geographically isolated HRTV lineages that diverged over the last several hundred years. Supporting this, there is evidence for geographic population structure within *A. americanum* ticks between different regions of the U.S.(48) Overall, our phylodynamic results for HRTV mirror what has been reported for the closely related Dabie bandavirus (DBV) in Asia. Our estimated evolutionary rate for HRTV, 1.1-1.8 x10^-4^ s/s/y, is very similar to the DBV literature, where studies report rates ranging from 1.07 x10^-4^ to 1.09 x10^-3^ s/s/y across the three segments, with most estimates around 2 x10^-4^.(34–36, 49, 50) Similar to our results, many DBV studies have shown distinct genotypes clustering by geographic region.(49, 51–53) Interestingly, however, several recent studies also report mixing of DBV genotypes across different countries in Asia,(7, 36) which is thought to be due to migratory bird movement carrying the DBV vector, *Haemaphysalis longicornis*, across longer distances.(54)

Despite the small number of HRTV sequences, our phylogenetic analysis suggests the possibility of reassortment between genome segments, a common mechanism of increasing genetic diversity in bunyaviruses and other segmented viruses.(55) Reassortment has been frequently described for DBV, allowing new genotypes to emerge after co-infection events.(7, 8, 56, 57) This is important because reassorted viruses have the ability to overcome host barriers and have the potential to generate strains of higher pathogenicity as seen in influenza virus.(55, 58) Because reassortment requires multiple viruses to infect the same cell, it would be expected to occur less frequently in viruses with low prevalence. Thus, our findings are surprising in light of the reported low HRTV prevalences in our study and others, underscoring the need for expanded surveillance and genetic studies of HRTV throughout the U.S.

Our observation of high HRTV genetic conservation at a focal site across multiple years suggests several possible hypotheses about the enzootic cycle of HRTV, which remains largely unknown.(3) One possibility is that closely related viruses may be sustained within the population through transovarial transmission (also referred to as vertical transmission), as has been documented in experimental studies of HRTV.(59) In this scenario, infected female *A. americanum* would lay eggs at the site, passing on virus to their larvae, which would feed on small-range hosts and remain within the site to molt and be captured as nymphs the following year. Vertical transmission typically augments other transmission mechanisms and has not been proposed as a sole mechanism for persistence of other tick-borne viruses.(60) Testing larval populations for HRTV is an important next step in investigating this mechanism.

A second potential mechanism for HRTV focality involves propagation of virus through cofeeding transmission, which has also been documented experimentally for HRTV(59) and is important in other tick-borne viruses such as tick-borne encephalitis virus (TBEV).(61) In this case, infected *A. americanum* feed next to uninfected ticks, leading to HRTV transmission through infection of local skin and leukocytes. Co-feeding transmission provides a role for hosts that are not competent for viremic infection themselves, as demonstrated for *Borrelia* species that feed on hosts whose innate complement systems prevent systemic infection.(62) Cofeeding on hosts with limited range could explain the persistence of highly related viruses across multiple years.

Finally, HRTV could propagate through horizontal transmission by ticks feeding on infected reservoir hosts. Serosurveys and experimental infections investigating potential HRTV reservoir hosts have not yet identified any candidates.(63, 64) Evidence of seropositivity has been found in multiple wildlife species with white-tailed deer, raccoons, horses, and dogs showing the highest rates of neutralizing antibodies across studies.(3, 16, 65) However, experimental infection of multiple wildlife hosts including white-tailed deer, raccoons, goats, chickens, rabbits, hamsters, and C57BL/6 mice have failed to produce detectable viremia, except in an interferon receptor-deficient mouse model.(63) Together, this information suggests that these animals may be incidental hosts, and there is not yet evidence for a viremic reservoir host to support horizontal transmission of HRTV. If there is a role for horizontal or cofeeding transmission in the HRTV enzootic cycle, our HRTV phylogenetic analysis suggests the importance of geographically restricted hosts. Our host bloodmeal analysis provides some possibilities, as *A. americanum* ticks at our focal site in central Georgia fed on deer and rabbits (which do not support HRTV viremia in experimental infection studies), as well as squirrels/chipmunks and birds (which have not been evaluated in experimental infection studies). While we did not observe a clear difference between HRTV-positive and negative samples, this analysis was limited by our analysis of tick pools rather than individual ticks, and it is possible that yet-untested hosts are important.

In conclusion, through extensive field work and the use of a novel multiplex amplicon sequencing assay, we have gained important insight into HRTV diversity and evolution. Key findings are that HRTV sequences from a focal site in GA were highly related across time; that there is strong geographic clustering of HRTV across the U.S. with this attribute being more important than host or year; and that all available HRTV sequences share a common ancestor within the recent past, suggesting that contemporary strains of HRTV emerged within the last 300 years. Our work offers potential insights into the enzootic cycle of HRTV and provides a foundation for further investigation.

## Acknowledgements

We thank undergraduate and graduate students from the Environmental Sciences Department at Emory University and Rollins School of Public Health for assistance with field work. We additionally thank Yamila Romer, Kayla Adcock and Daniel Mead for their initial work on Heartland virus detection and isolation from ticks in Georgia.

## Author Contributions

SB, GVP, and AP conceived of the study. Field collections were conducted by SB, NS, AL, LA, EF, TW, SC, IR, and MS. Lab processing, amplicon sequencing design and sequencing was completed by SB, NS, EO, MS and HD. Phylogenetic analyses were conducted by SB, TF, and AP. Host bloodmeal analysis was conducted by HKG and SRT. All authors read, edited, and approved the final manuscript.

