## Supplement for "Focal persistence and phylodynamics of Heartland virus in Georgia"

**Supplement File. Focal persistence and phylodynamics of Heartland virus in Georgia**

**Supplemental Table S1. Extraction Procedure**

| | |
|---|---|
| Step 1 | In 1.5 mL tubes labeled by sample number, add 560 uL of Buffer AVL |
| Step 2 | Add 5.6. uL of linear acrylamide* into each tube |
| Step 3 | Pipette 140 uL of each sample into corresponding 1.5mL conical tube and mix by vortex for 15s and centrifuge |
| Step 4 | Incubate at room temp for 10 min (the virus is now inactivated) |
| Step 5 | Add 560uL of 100% ethanol to sample and mix by vortex for 15s. Briefly centrifuge to remove drops from inside of lid |
| Step 6 | Add 630uL of sample to column (labeled) and centrifuge for 1 min @ 8000rpm. Discard collection tube. Place column in clean collection tube. |
| Step 7 | Add remainder of sample to column and centrifuge for 1 min @ 8000rpm. Discard collection tube. Place column in clean collection tube. |
| Step 8 | Add 500uL of Buffer AW1 to column. Centrifuge for 1 min @ 8000rpm. Discard collection tube. Place column in clean collection tube. |
| Step 9 | Add 500uL of Buffer AW2 to column. Centrifuge for 3 min @ 14,000rpm. Discard collection tube. Place column in clean collection tube. |
| Step 10 | Centrifuge again for 1 min @ 14,000rpm (essential to remove excess ethanol from column) |
| Step 11 | Place column in a new (labeled) 1.5ml microcentrifuge tube. |
| Step 12 | Add 60ul of Buffer AVE to column at the center of the filter. Incubate at room temp for at least 1 min. Centrifuge for 1 min @ 8000rpm. |
| Step 13 | Aliquot 30 uL of the eluted total nucleic acid (TNA) into a sticker-labeled screw cap PCR tube (cryovial) for long-term storage and store the remainder in the 1.5 microcentrifuge tube in the RT-qPCR box in the -80 |

*Utilized linear acrylamide as added nucleic acid backbone instead of kit provided carrier RNA to prevent overwhelming background during metagenomic sequencing

**Supplemental Table S2. Heartland RT-qPCR Assay Information**

| Primer and Probe information | |
|---|---|
| **Name** | **Sequence (5’-3’)** |
| HRTV S1 Forward | TGCAGGCTGCTCATTTATTC |
| HRTV S1 Reverse | CCTGTGGAAGAAACCTCTCC |
| HRTV S1 Probe | CCTGACCTGTCTCGACTGCCCA |

| Reaction Concentrations (Kit: Ag-Path 1 Step ID Kit) | |
|---|---|
| **Reagent** | **Amount per well (uL)** |
| Water | 3 |
| 2X QuantiTect Mix | 12.5 |
| HRTV Forward primer | 1 |
| HRTV Reverse primer | 1 |
| HRTV Probe | 1.5 |
| 25X RT-PCR Enzyme Mix | 1 |
| Sample | 5 |
| Total | 25 |

| Cycling Conditions | |
|---|---|
| 45°C | 10 min |
| 95°C | 10 min |
| 45 cycles: | |
| 95°C | 15 sec |
| 60°C | 45 sec |

**Supplemental Table S3. Tick Actin RT-qPCR Assay Information**

| Name | Sequence (5'-3') |
|---|---|
| Tick Actin Forward | GGACAGCTACGTGGGCGACGAGG |
| Tick Actin Reverse | CGATTTCACGCTCAGCCGTGGTGG |

| Reaction Concentrations | |
|---|---|
| Kit: Applied Biosystems Power SYBR Green RNA-to-Ct 1-step PCR | |
| **Reagent** | **Amount per well (uL)** |
| Water | 4.04 |
| Power SYBR Mix | 10 |
| Tick Actin Fwd primer | 0.4 |
| Tick Actin Rev primer | 0.4 |
| 125x RT Enzyme Mix | 0.16 |
| Sample | 5 |
| Total | 20 |

| Cycling Conditions | |
|---|---|
| 48°C | 30 min |
| 95°C | 10 min |
| 45 cycles: | |
| 95°C | 15 sec |
| 60°C | 45 sec |
| Melt Curve | |
| 95°C | 15 sec |
| 60°C | 15 sec |
| 95°C | 15 sec |

### Supplemental Table S4. Custom primer sequences for the HRTV amplicon primer set.

| L Primer Name | Primer Sequence | M Primer Name | Primer Sequence | S Primer Name | Primer Sequence |
|---|---|---|---|---|---|
| HRTV_L_1_FWD | TCCAGATGAATTTAGAAGCTCTTTGC | HRTV_M_1_FWD | ATCCACTGAGATGATTGTCCCG | HRTV_S_1_FWD | ACACAAAGAACCCCTTGAATTATCAA |
| HRTV_L_1_REV | CCAGAGAAAGTGAAGTCATGATTTATCT | HRTV_M_1_REV | GGAAACACTTGGTCTTCCCTTC | HRTV_S_1_REV | CAAGAAGTTGCTCAACAGCTGG |
| HRTV_L_2_FWD | GCTAGGATCCTCGATCAATGCA | HRTV_M_2_FWD | TTGGACATGTCACAAGCTGGTT | HRTV_S_2_FWD | CCTTCACCAATACTGTCTGGCT |
| HRTV_L_2_REV | CGATAAGCAGCCTCAAGTCCTC | HRTV_M_2_REV | CCCAGCCTTTGTAACCGTTGTA | HRTV_S_2_REV | TCCTGTGGAAGAAACCTCTCCA |
| HRTV_L_3_FWD | CCACACAAGACTAGATGGAACAGT | HRTV_M_3_FWD | GGCTGGAGACACAGACATGATT | HRTV_S_3_FWD | CTTCTCCCTCAAGAACACCTGG |
| HRTV_L_3_REV | TTTGCCACACAGAACCTGAACA | HRTV_M_3_REV | GGGAACTCACTTTGACTCAAGCT | HRTV_S_3_REV | GAGACCCCTAACACGCAAAAGT |
| HRTV_L_4_FWD | TGTTAGTGCATCTGGTGTTCTCA | HRTV_M_4_FWD | GAAGTGGTCTGTCAGAGAGGGA | HRTV_S_4_FWD | CAATAATGGACGTGGCTCTCCT |
| HRTV_L_4_REV | TTCTGAGCATCTCAATGTCAGAGT | HRTV_M_4_REV | GACCCACCTTGAAGCACACAAA | HRTV_S_4_REV | GGGCAAGTCAGACAGGTACAAG |
| HRTV_L_5_FWD | AGCTTTCTTCACGCTTTTTGACT | HRTV_M_5_FWD | AGGTACTGATGAGAGAACACAAAACT | HRTV_S_5_FWD | ACTTTGCCATCTGGAGAGACAA |
| HRTV_L_5_REV | TCCATGAGAGCTCTAGTTTTATCT | HRTV_M_5_REV | TCTCACAGAGACGCCTCCATAA | HRTV_S_5_REV | CTAAGGGGAGCTGGAAAGTGCT |
| HRTV_L_6_FWD | AGTGGCATGAGCACAAAAGAGA | HRTV_M_6_FWD | TGCCCGTATGCAAGATAATCAAGA | HRTV_S_6_FWD | ACATTATCTGAATAAATGCCCTCTCCA |
| HRTV_L_6_REV | CCAGGAAGCATCTTGACCACAA | HRTV_M_6_REV | GCAGATGCACAAAAATGTGAAGC | HRTV_S_6_REV | AATGCCACCAGGATCAGCTG |
| HRTV_L_7_FWD | GTCAAAACGGACACGGGATCAG | HRTV_M_7_FWD | GGGAGGAGTCAAAATTGTAACGC | HRTV_S_7_FWD | CTGCTGCAGCCTTCACTGAG |
| HRTV_L_7_REV | GCCCTCGATGCCTTGTAATGAA | HRTV_M_7_REV | GTGGGTTTCTCAGGAACTCCTT | HRTV_S_7_REV | GGACTTCATGGATGCCTATTCCC |
| HRTV_L_8_FWD | TAAAATACCACAGGTTCCGGCC | HRTV_M_8_FWD | GGGCACTGCATATTCCCAGATG | HRTV_S_8_FWD | AGGGTCTCTGAATTTGGCATACA |
| HRTV_L_8_REV | TAGCCTTCTCCAACAACAGGTT | HRTV_M_8_REV | ACCCCTTCTTTGACAGTTTGGC | HRTV_S_8_REV | GCAGCAGCAATCAAGGAGTAC |
| HRTV_L_9_FWD | TGAACAACCTGTTAAATGACACATCT | HRTV_M_9_FWD | TGTTTGGGGGACATGGGTTTTT | HRTV_S_9_FWD | TCATTTCCAGTGGATAACCTTGGA |
| HRTV_L_9_REV | TGGCCAGTGCATTAATTTTTTGA | HRTV_M_9_REV | TCACACCCAGAGCAAATTGAAGT | HRTV_S_9_REV | ATTTGCTCTCACCAGAGGGAA |
| HRTV_L_10_FWD | TAGCTCAGGAATTGGCAAGTGC | HRTV_M_10_FWD | GACAAGGTTATGGCCCAAGAGG | HRTV_S_10_FWD | CATCCTCTCAGCGCCTTTCTTA |
| HRTV_L_10_REV | AGATTTGTCAGCTTGCACGTCT | HRTV_M_10_REV | TGATCTTGAGGCACTTAGAGTCTG | HRTV_S_10_REV | GCATGACTGACTGGTCTGCAAT |
| HRTV_L_11_FWD | TCACAGACACAATTGATGCGGG | HRTV_M_11_FWD | TTGCTCCCAGCTGTTAATCCA | | |
| HRTV_L_11_REV | CGTGAGAGTGACAAGCTCTTCT | HRTV_M_11_REV | CAGCCCTGTCCATCCTGTTT | | |
| HRTV_L_12_FWD | CTCAGGGAACAAGTGGCAATCA | HRTV_M_12_FWD | GTGACTGTCAATCTGGGTGTCC | | |
| HRTV_L_12_REV | CATCTCTTGCCACAATGGGGAA | HRTV_M_12_REV | GTCTTAACCTCTCCTGATGGCA | | |
| HRTV_L_13_FWD | GCGAACAAAGCTTCAAGTGTTTT | HRTV_M_13_FWD | AATCGTGTCTGGAAGGTGTCAC | | |
| HRTV_L_13_REV | TCTCAGCCTCAATCTCAACAATCT | HRTV_M_13_REV | CACATTTCCCATTATGGCGTGC | | |
| HRTV_L_14_FWD | AAAAACAAAGAAGAGTCGACCGAG | HRTV_M_14_FWD | TGAAAGAGATGGGAACTGGGAT | | |
| HRTV_L_14_REV | TGAGCCCAGTTCTTTAAGGACTC | HRTV_M_14_REV | ACCCCAGAAATTGTTGCCGG | | |
| HRTV_L_15_FWD | GCATGCATATCTCTTGAGAAGGAGA | HRTV_M_15_FWD | AGGGTTGATGATGCTGTGTGTT | | |
| HRTV_L_15_REV | TGACCTGCAGCATCAACATACC | HRTV_M_15_REV | ACCAGAGACGTTTGTTATCTCACC | | |
| HRTV_L_16_FWD | CATAGGTCCAAACTCCTAGAAAAGATG | HRTV_M_16_FWD | TGTCATTCAGAGGGATGAGGCT | | |
| HRTV_L_16_REV | GGGTTAGCTATTGTTTCATGTGGG | HRTV_M_16_REV | CAGTTTTGATGAGTGGCCACCA | | |
| HRTV_L_17_FWD | TGCAAGGCTTGTCCAATTTGGA | HRTV_M_17_FWD | AGGCCTCACATGAGCTTTGATA | | |
| HRTV_L_17_REV | AACCTATGGAACTTTGCTGGCA | HRTV_M_17_REV | GCCAGCAGAACCCCTATAATCC | | |
| HRTV_L_18_FWD | TGGAGTCAGGGTCATTATACAACC | HRTV_M_18_FWD | TAGACTGGCTCAACGCACTGTT | | |
| HRTV_L_18_REV | ATGTGGCTCCTTGAGTCATCCA | HRTV_M_18_REV | ACCCACCCATTATCATTCATGCC | | |
| HRTV_L_19_FWD | TCAGAAAACTGATGATGACTTCAGGA | | | | |
| HRTV_L_19_REV | TCAGAGCCTTCTAAAACATCTATAGCT | | | | |
| HRTV_L_20_FWD | GCGTCCAAAGTTTTTACAAGGCA | | | | |
| HRTV_L_20_REV | ATAAGGTATTTACTGTTGACTTCTCAGAG | | | | |
| HRTV_L_21_FWD | ACAGCCAACTTGCTATACTCAGT | | | | |
| HRTV_L_21_REV | ATGATGAACCCCCTTCAAGACA | | | | |
| HRTV_L_22_FWD | TCTGATTCAGAAGCCCTGGCAA | | | | |
| HRTV_L_22_REV | TAAATCCAGCTCCCCCTGCAAA | | | | |
| HRTV_L_23_FWD | GGGACATTTGTAGGGATGCTGA | | | | |
| HRTV_L_23_REV | CCAGTAGATCATTACAGAGTGGCTT | | | | |
| HRTV_L_24_FWD | CCAAGGGGGATGCAGATTACAG | | | | |
| HRTV_L_24_REV | CTGAAGCTGGAAGTGACACCTG | | | | |
| HRTV_L_25_FWD | ATAGGAGGCCTGAGAACAAGC | | | | |
| HRTV_L_25_REV | GCCTTCCTTGATTCCTCTCAGC | | | | |
| HRTV_L_26_FWD | TACACTGCATCAATCCACGGTA | | | | |
| HRTV_L_26_REV | AACCACATCTCTGCAATGAGGT | | | | |
| HRTV_L_27_FWD | AAGTTTGTTGTGAGAGAAAGGAACA | | | | |
| HRTV_L_27_REV | TGTGCAATGAAATTCCTGAATTGAAC | | | | |
| HRTV_L_28_FWD | GCACTGACCATAAAGAAACACTGGA | | | | |
| HRTV_L_28_REV | TTTGCTCATCAGTGTAAGGGCC | | | | |
| HRTV_L_29_FWD | CCCAGGTTTCATTTTGGACTCCA | | | | |
| HRTV_L_29_REV | ATTGACCCTCCTCTCTGCTCAA | | | | |
| HRTV_L_30_FWD | GCTGCTGAGGTTGTGAAGAGAT | | | | |
| HRTV_L_30_REV | CCAGTTCGATGAACCATCACCA | | | | |
| HRTV_L_31_FWD | AAGGGGAAAGGTGTTTGGTCAG | | | | |
| HRTV_L_31_REV | CACCCTTCTACCCTTGAAGCTG | | | | |
| HRTV_L_32_FWD | TAACATCAAGCTTTCGGGGTCA | | | | |
| HRTV_L_32_REV | GGATGCTGCAGACTCTGAGATG | | | | |
| HRTV_L_33_FWD | CTTCAAGAGGACCGAGTCATGA | | | | |
| HRTV_L_33_REV | CCTTGAAAATGTTCCCCATGGC | | | | |
| HRTV_L_34_FWD | GGAATCAGGCAGCAAGAGTTCT | | | | |
| HRTV_L_34_REV | TCAATGTCTAGGTCCCCTTCCA | | | | |
| HRTV_L_35_FWD | ACAAGTCAGGAGTTGGTGGATA | | | | |
| HRTV_L_35_REV | CCTGCAACCATCTTTTCCAAGG | | | | |
| HRTV_L_36_FWD | TGGTGATACTCAGTCAGTCAGGA | | | | |

**Supplemental Table S5: Pairwise genetic distances between all Heartland virus sequences in the U.S.**

| | GA23_T_2019 | GA26_T_2019 | GA100_T_2022 | GA404_T_2021 | GA187_T_2022 | GA504_T_2019 | GA056F_T_2023 | GA432R_T_2023 | GA642T_T_2023 | H_KY_2017 | H_TN_2013 | s1782_T_NY_2018 | s1820_T_NY_2018 | NC02449X_H_MO_2009 | H_MO_2017 | H_OK_2017 | JX00584X_H_MO_2009 | sM1266_T_MO_2012 | M1275_T_MO_2012 | T_KS_2015 |
|---|---|---|---|---|---|---|---|---|---|---|---|---|---|---|---|---|---|---|---|---|
| GA26_T_2019 | 99.97 | | | | | | | | | | | | | | | | | | | |
| GA100_T_2022 | 99.91 | 99.93 | | | | | | | | | | | | | | | | | | |
| GA404_T_2021 | 99.92 | 99.93 | 99.98 | | | | | | | | | | | | | | | | | |
| GA187_T_2022 | 99.9 | 99.92 | 99.99 | 99.97 | | | | | | | | | | | | | | | | |
| GA504_T_2019 | 99.78 | 99.79 | 99.78 | 99.8 | 99.77 | | | | | | | | | | | | | | | |
| GA056F_T_2023 | 99.9 | 99.91 | 99.93 | 99.93 | 99.92 | 99.75 | | | | | | | | | | | | | | |
| GA432R_T_2023 | 99.89 | 99.9 | 99.92 | 99.91 | 99.91 | 99.74 | 99.93 | | | | | | | | | | | | | |
| GA642T_T_2023 | 99.55 | 99.58 | 99.56 | 99.56 | 99.55 | 99.52 | 99.55 | 99.56 | | | | | | | | | | | | |
| H_KY_2017 | 97.27 | 97.28 | 97.32 | 97.32 | 97.31 | 97.24 | 97.29 | 97.28 | 97.31 | | | | | | | | | | | |
| H_TN_2013 | 98.16 | 98.17 | 98.2 | 98.2 | 98.19 | 98.13 | 98.17 | 98.17 | 98.24 | 98.62 | | | | | | | | | | |
| s1782_T_NY_2018 | 97.73 | 97.74 | 97.75 | 97.75 | 97.74 | 97.68 | 97.78 | 97.75 | 97.78 | 96.87 | 97.73 | | | | | | | | | |
| s1820_T_NY_2018 | 97.67 | 97.68 | 97.68 | 97.68 | 97.68 | 97.62 | 97.71 | 97.68 | 97.71 | 96.82 | 97.68 | 99.88 | | | | | | | | |
| NC02449X_H_MO_2009 | 97.82 | 97.83 | 97.84 | 97.86 | 97.84 | 97.79 | 97.85 | 97.83 | 97.78 | 97.02 | 97.88 | 98.36 | 98.24 | | | | | | | |
| H_MO_2017 | 97.79 | 97.8 | 97.83 | 97.85 | 97.82 | 97.74 | 97.82 | 97.79 | 97.77 | 97.02 | 97.89 | 97.45 | 97.38 | 97.54 | | | | | | |
| H_OK_2017 | 95.11 | 95.14 | 95.15 | 95.15 | 95.14 | 95.07 | 95.16 | 95.13 | 95.13 | 94.89 | 95.14 | 95.47 | 95.39 | 95.48 | 94.86 | | | | | |
| JX00584X_H_MO_2009 | 96.91 | 96.94 | 96.95 | 96.95 | 96.94 | 96.86 | 96.96 | 96.93 | 96.93 | 96.12 | 96.96 | 97.37 | 97.25 | 97.34 | 96.69 | 97.46 | | | | |
| sM1266_T_MO_2012 | 94.41 | 94.44 | 94.45 | 94.45 | 94.44 | 94.36 | 94.46 | 94.43 | 94.43 | 94.12 | 94.46 | 94.82 | 94.74 | 94.8 | 94.14 | 97.43 | 97.36 | | | |
| M1275_T_MO_2012 | 96.82 | 96.85 | 96.85 | 96.85 | 96.85 | 96.77 | 96.86 | 96.84 | 96.84 | 96.38 | 96.86 | 96.88 | 96.81 | 96.85 | 96.55 | 97.38 | 99.47 | 97.34 | | |
| T_KS_2015 | 94.43 | 94.46 | 94.47 | 94.47 | 94.46 | 94.4 | 94.48 | 94.45 | 94.47 | 94.29 | 94.5 | 94.82 | 94.74 | 94.8 | 94.16 | 97.63 | 96.81 | 97.72 | 96.65 | |
| H_KY_2015 | 95.93 | 95.93 | 95.94 | 95.96 | 95.93 | 95.91 | 95.96 | 95.93 | 95.96 | 95.21 | 96.02 | 95.74 | 95.63 | 95.8 | 95.94 | 93.55 | 95.28 | 92.76 | 95.12 | 92.84 |

Percent nucleotide identity is displayed across the concatenated genome of the three segments excluding gaps in the sequences generated. Sequences are named by the state they were collected in, host of sample, year and if that name was still not unique, the isolate number assigned in GenBank. Background color gradient represents nucleotide identity with darker colors represent higher nucleotide identity shared between sequences.

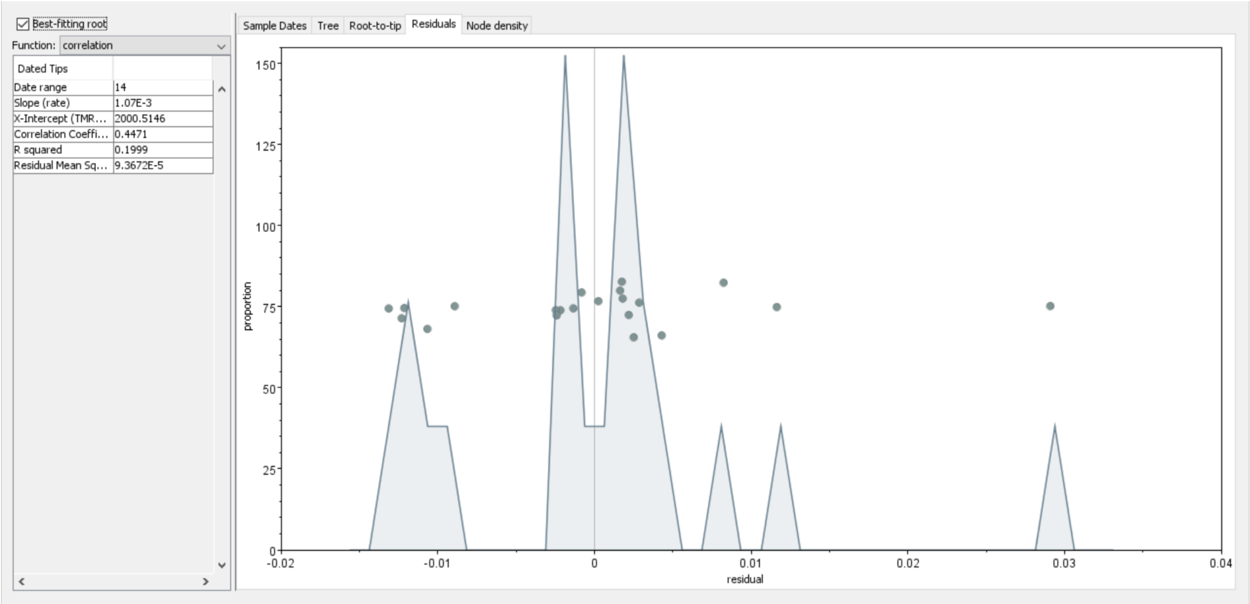


**Supplemental Figure S1. Temporal residual histogram of HRTV L segments** containing OK480062.1_H_L_KY_2015 (far right point) demonstrating unbalanced residual of ~0.03 and was therefore removed from evolutionary rate analysis.

**Supplemental Text S1. HRTV amplicon sequencing protocol**

#### Custom Primer Design

Custom primers were designed using an alignment of 4 (L) or 5 (M/S) complete (>95%) Heartland (HRTV) reference genome sequences. Alignments of the Large, Medium, and Small (L, M, S) segments of the HRTV genome were loaded into PrimalScheme v.1.4.1 to generate 36, 18, and 10 primer pairs for the L, M, and S segments respectively (See Supplementary Table S4). Each primer pair was designed to amplify a fragment of approximately 250 base pairs (bp) with a melting temperature 60ºC ±3C. The primers created by PrimalScheme were then mapped to the individual reference sequences in order to confirm proper amplicon length and evaluate mismatches. Primers with multiple (>3) mismatches or a mismatch within 3bp of the 3' end were modified in Geneious Prime (www.geneious.com) while still ensuring a melting temperature of 60ºC ±3C. Primer-index fusions were generated as described by Glenn et al.[1] by adding the forward (5'TCGTCGGCAGCGTCAGATGTGTATAAGAGACAG3') or reverse (5'GTCTCGTGGGCTCGGAGATGTGTATAAGAGACAG3') Nextera iNext universal adaptor sequence to the 3' end of each custom HRTV primer. Fusion primers were synthesized by Integrated DNA Technologies (IDT, Iowa, USA) and pooled to equimolar concentration at 10mM. Two primer pools were created, with even and odd-numbered primer pairs separated in order to prevent undesired primer pairing during multiplex PCR. Pilot experiments were conducted using the previously-sequenced HRTV-positive tick lysate samples, and primer concentrations were adjusted to ensure even coverage across the genome.

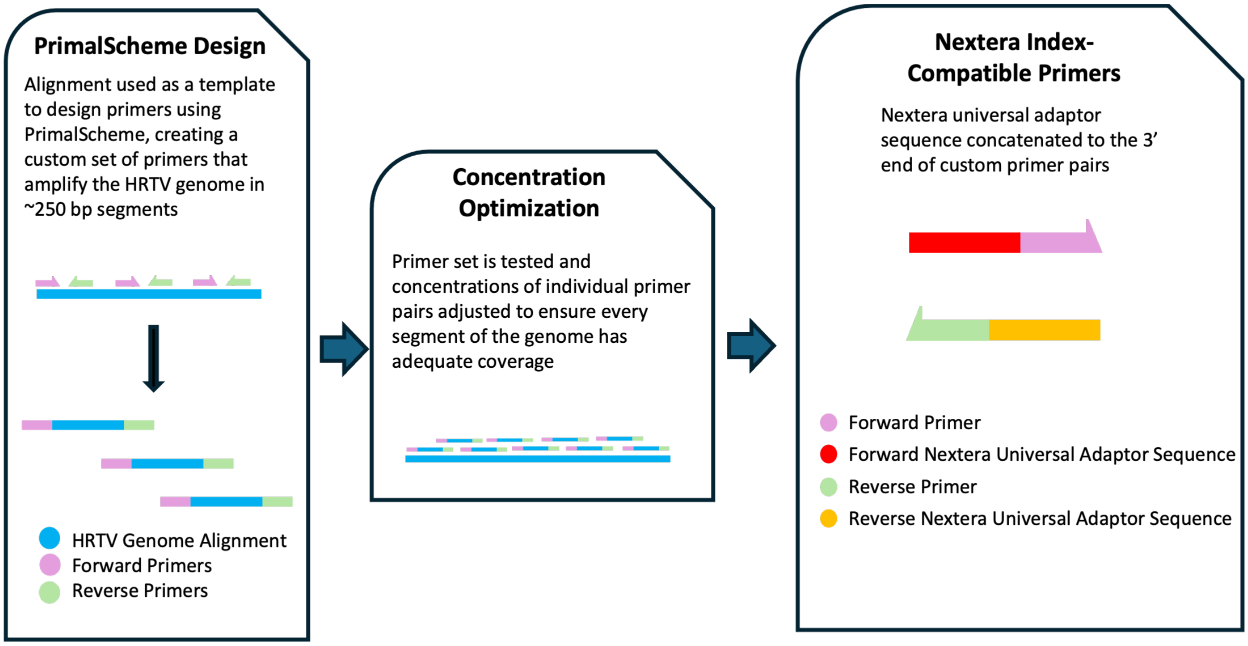

**Library Construction and Sequencing**

1. **DNase Treatment**

Extracted RNA underwent heat-labile DNase treatment (ArcticZymes, Tromso, Norway) by adding 12µl of extracted RNA to a DNase reaction mix consisting of 1.2 µl DNase Buffer and 1 µl of DNase enzyme and incubated at 37°C for 10 minutes, then 58°C for 5 minutes.

2. **Single-stranded cDNA Synthesis**

Single- stranded cDNA was generated from the DNA-depleted RNA using the Superscript IV (SSIV) First-Strand Synthesis System (Fisher). First, 5.28 µl RNA was primed by mixing with 4.72 µl (insert concentration here) random hexamers and heating to 65ºC for five minutes. Then, the SSIV reaction mix consisting of 4 µl 5X SSIV buffer, and 1 µl each of .1M DTT, 10mM dNTP mix, nuclease-free water, SUPERase-ln (20U/ µl), Superscript IV RT (9 µl total) was added to the primed sample. This mixture was incubated at 23°C, then 55°C, then finally 80°C for 10 minutes each.

3. **Multiplex PCR**

Single-stranded cDNA from each sample was aliquoted into two wells, with 3µl cDNA per well. To each well was added a reaction mix consisting of 12.5 µl Q5 2X Hot Start DNA Polymerase Mix (NEB), 7.5 µl nuclease-free water, with 10 µl of either even or odd 10mM primer pool. Samples were amplified by incubating at 98°C for 30 seconds, then 18 cycles of 95°C for 15 seconds and 65°C for 5 minutes (each cycle 5:15 minutes total), and finally held at a temperature of 4°C. This process generated the ~250bp HRTV amplicons with an affixed Nextera iNext universal adaptor sequence on each end. Even and odd pools for each sample were combined and purified by performing a .7X SPRI using Ampure XP beads (Beckmann-Coulter).

4. **Indexing Reaction**

Samples were indexed using Nextera indices, which contain a region complementary to the universal adaptor sequence as well as a unique region for identification of individual samples. A 5µl aliquot of each amplicon was added to a well with 12.5 µl Q5 2X Hot Start DNA Polymerase Mix (NEB), 5 µl nuclease-free water, and 1 µl dual-unique Nextera index pair. The following thermocycling conditions were used: 98°C for 30 seconds, 10 cycles of 95°C for 15 seconds, 60°C for 15 seconds, and 72°C for 30 seconds (each cycle 1 minute total), and finally 72°C for 5 minutes. This created HRTV amplicons with unique Nextera indices for each sample. The indexed amplicons were purified by performing another .7X SPRI cleanup.

5. **KAPA, Pooling, and Sequencing**

The indexed libraries were quantified using the KAPA Universal Complete Kit (Illumina) according to kit instructions. Samples were pooled to equimolar concentration, purified by performing a .8X SPRI and the pool concentration was quantified using KAPA. The pool was then loaded on a MiSeq 300bp v2 paired-end read kit and sequenced.
